## Supplementary figures for "Reference genome and resequencing of 305 accessions provide insights into spinach evolution, domestication and genetic basis of agronomic traits"

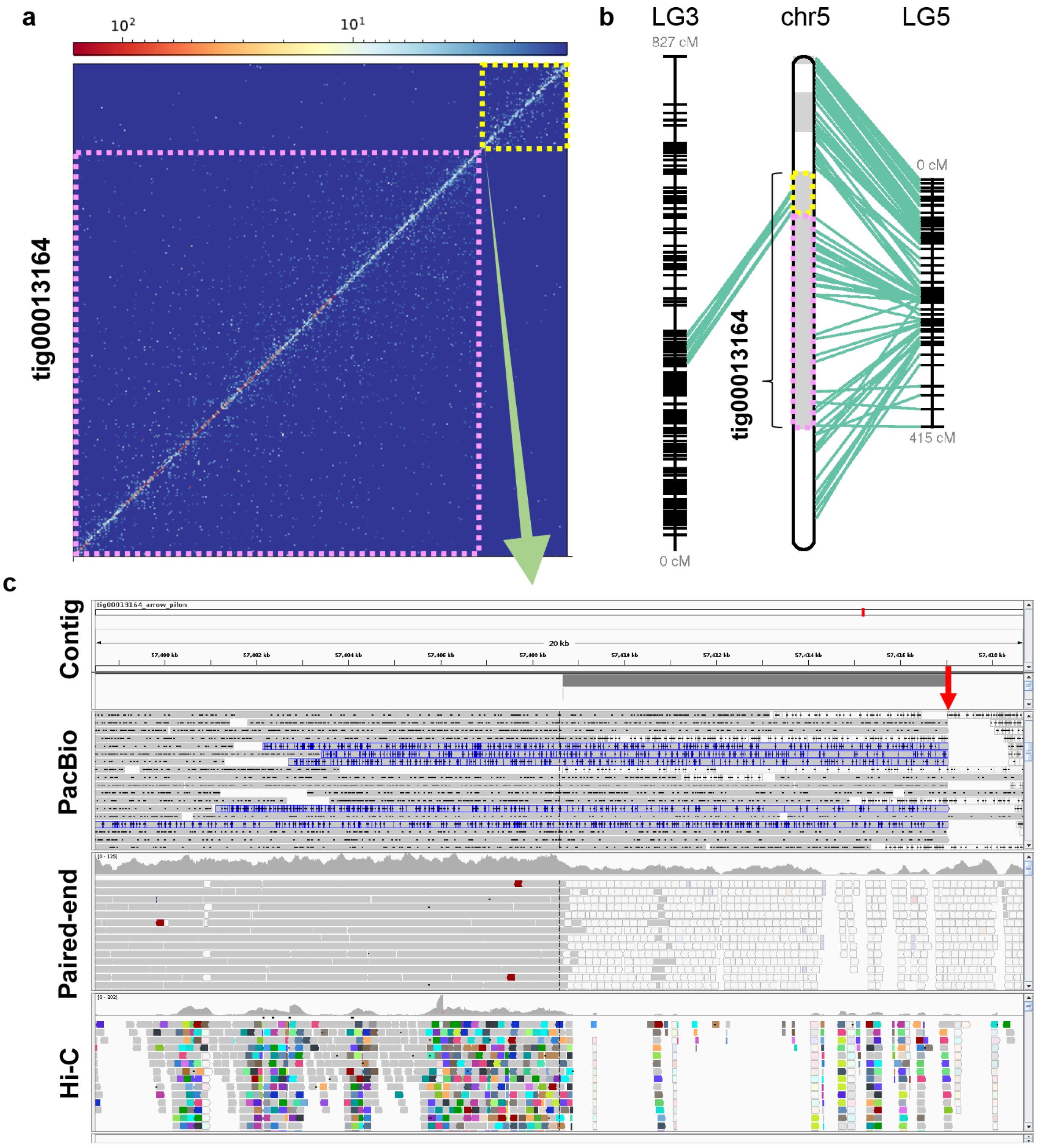


**Supplementary Fig. 1 Example of a misassembled contig in the initial PacBio long read assembly.** (**a**) Heatmap of Hi-C interactions of contig tig00013164, indicating a possible misjoining of two regions (marked with yellow and pink dashed-lines, respectively) indicated by very few Hi-C interactions. Color bar at the top represents the density of Hi-C interactions, which are indicated by number of links at the 50-kb resolution. The misassembly and the break point are supported by the evidence from genetic maps (**b**) and alignments of PacBio long reads, Illumina paired-end reads and Hi-C long contact reads (**c**). Most of the PacBio long reads were either ended or split (reads with blue border) at the suggested break point (pointed with red arrow).

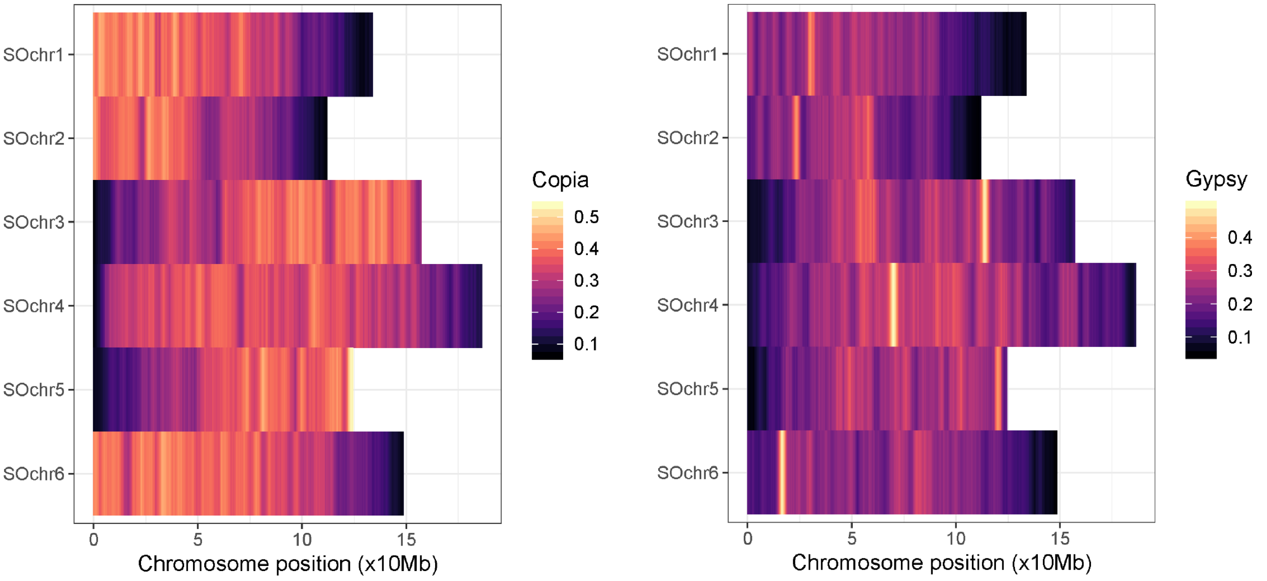


**Supplementary Fig. 2 Distribution of *Copia* and *Gypsy*-type LTR retrotransposons in the 'Monoe-Viroflay' genome.**

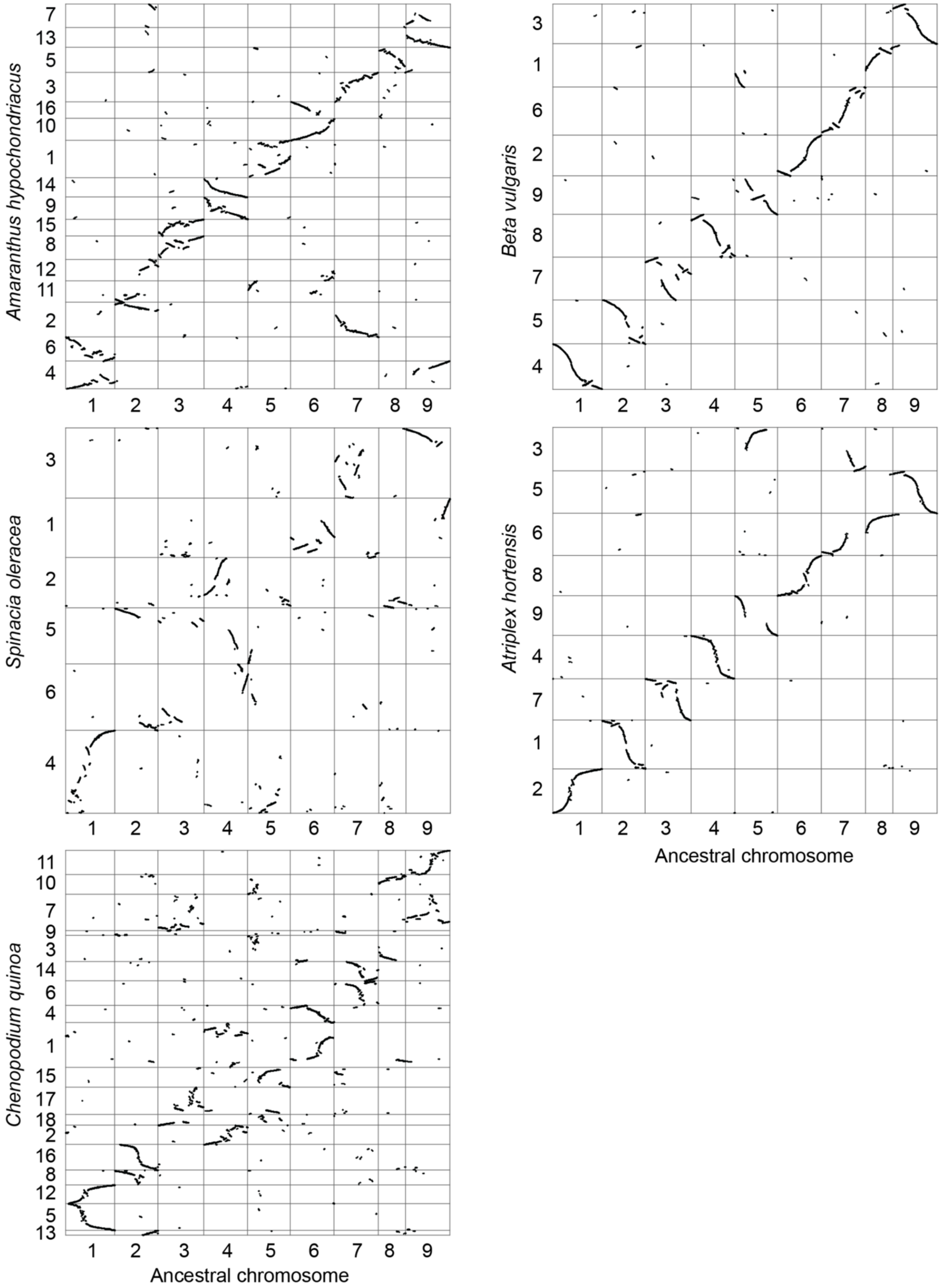


**Supplementary Fig. 3 Genome synteny between extant and ancestral Chenopodiaceae.**

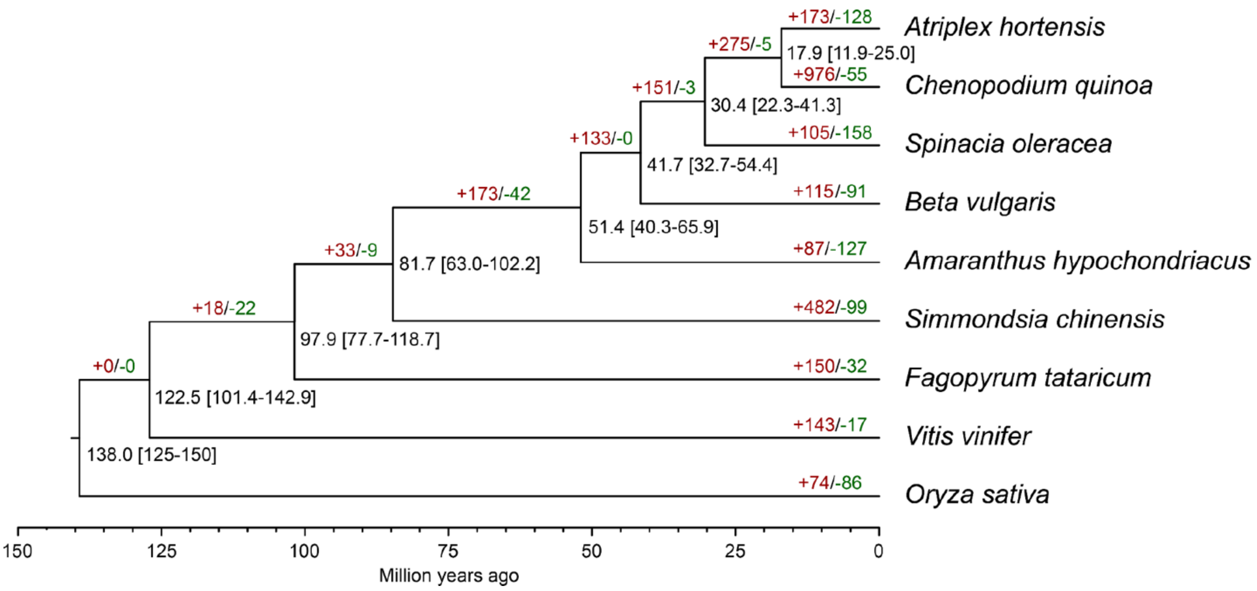


**Supplementary Fig. 4 Phylogeny and gene family evolution of selected species.** Black numbers around the branch of the tree represent the divergence time (million years ago) and the 95% highest posterior density range (in the bracket). Red and green numbers on the tree represent numbers of expanded (+) and contracted (-) gene families across the evolution of the species.

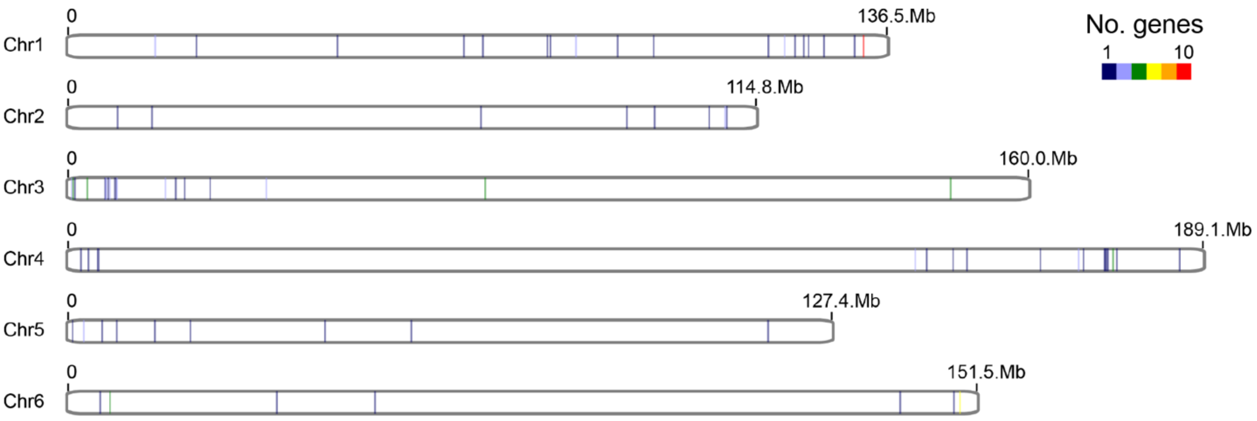


**Supplementary Fig. 5** Distribution of NBS-LRR genes in the 'Monoe-Viroflay' genome. The NBS-LRR gene clusters and singletons are plotted on each chromosome. The size of gene clusters is indicated by the color.

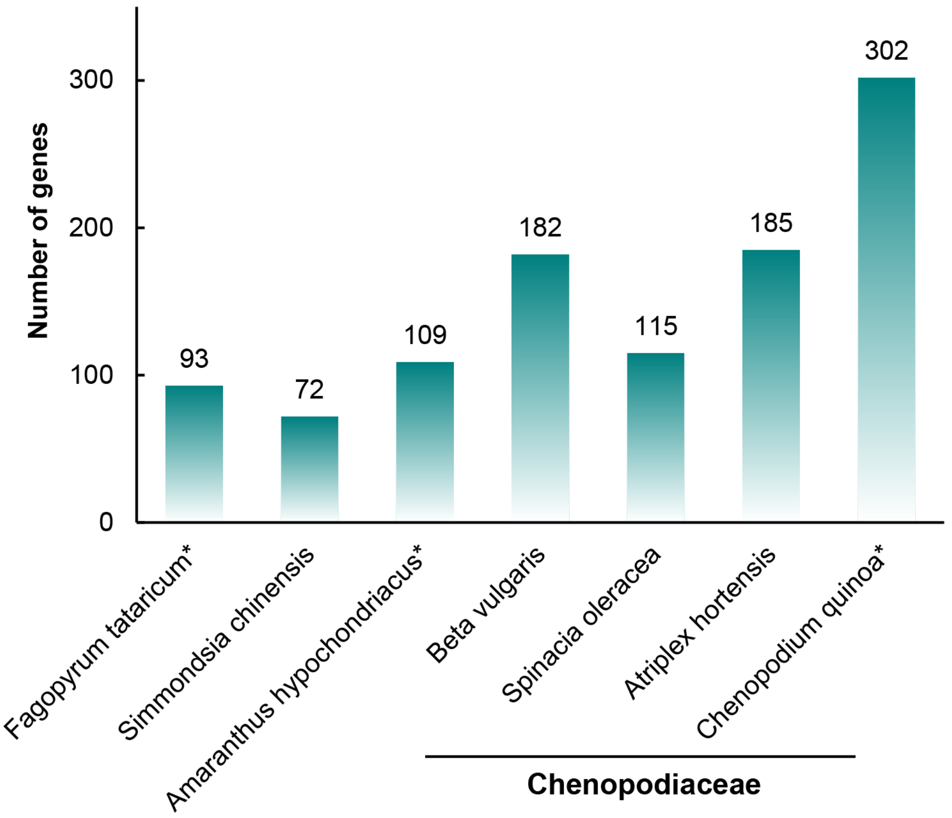


**Supplementary Fig. 6 Number of NBS-LRR genes in the selected species.**

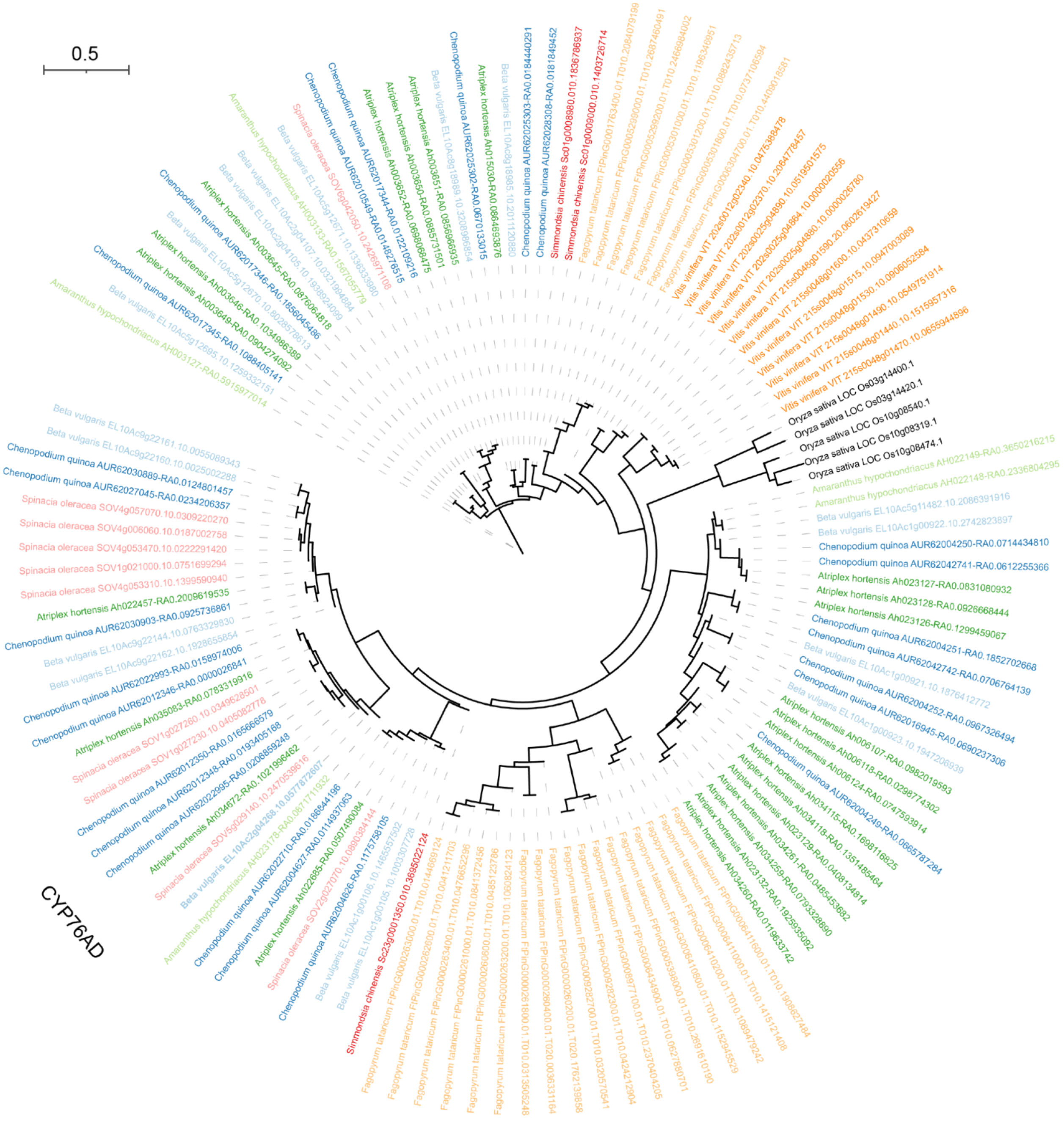


**Supplementary Fig. 7 Phylogeny of CYP76AD genes in the selected species.** Species are labeled in different color.

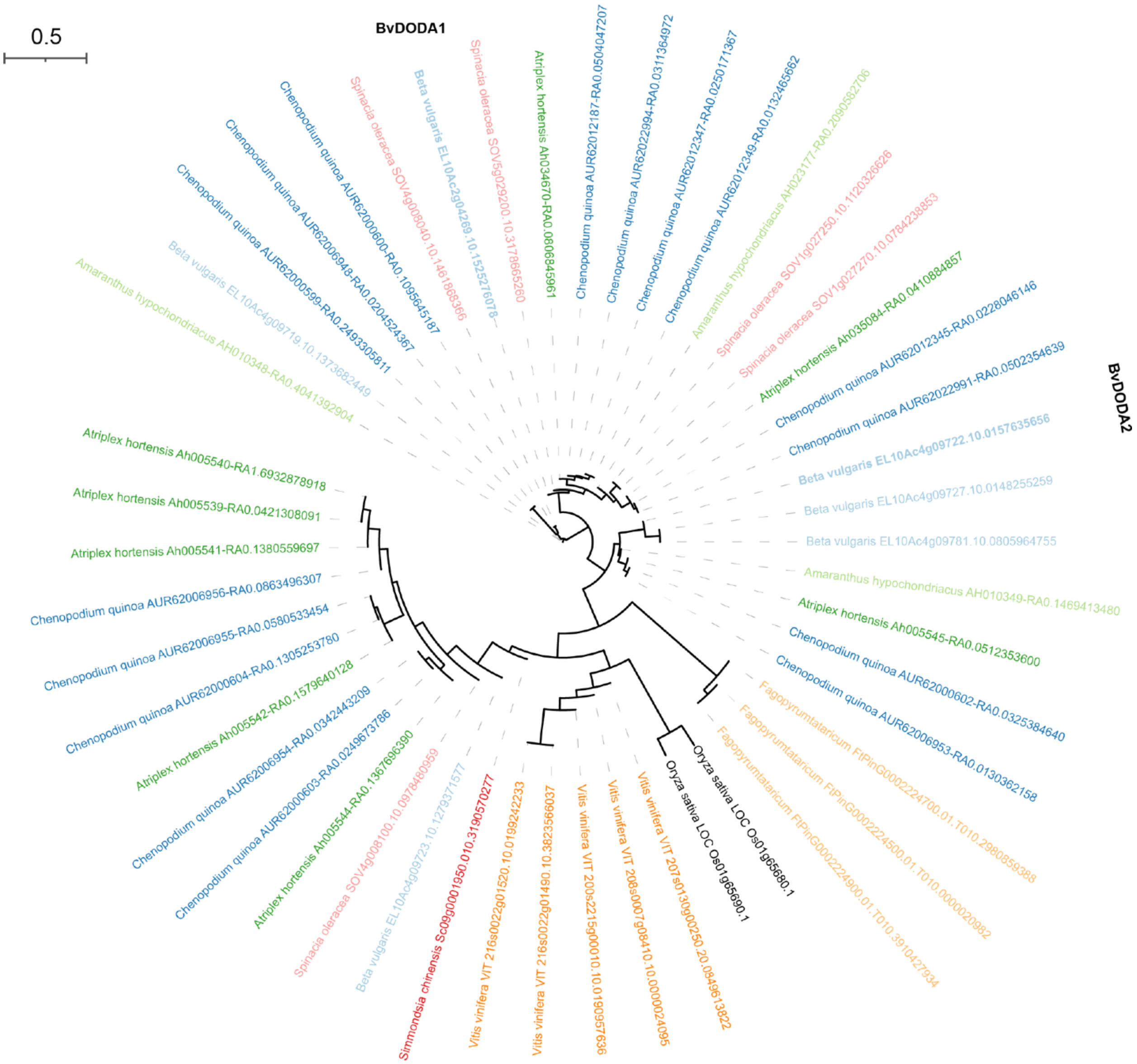


**Supplementary Fig. 8 Phylogeny of DODA genes in the selected species.** Species are labeled in different color.

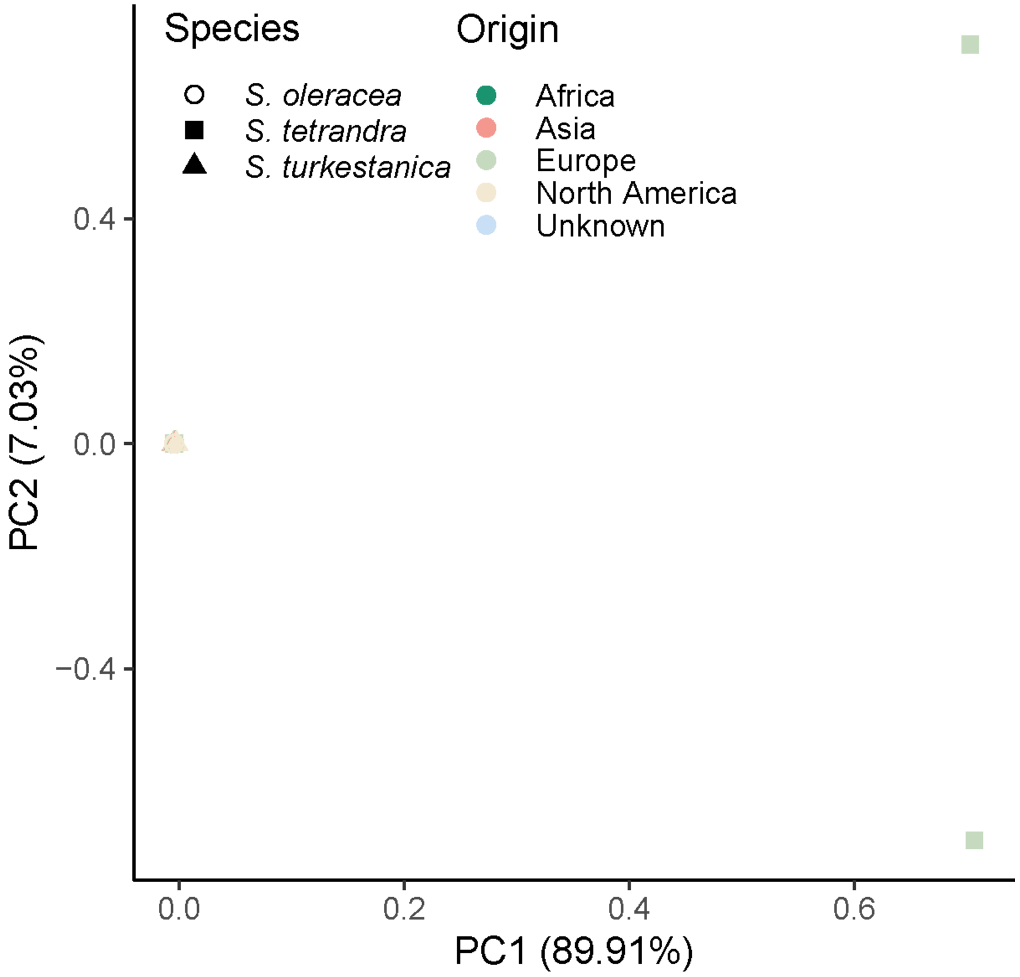


**Supplementary Fig. 9. Principal component analysis of *Spinacia* accessions using SNPs at fourfold degenerate sites.**

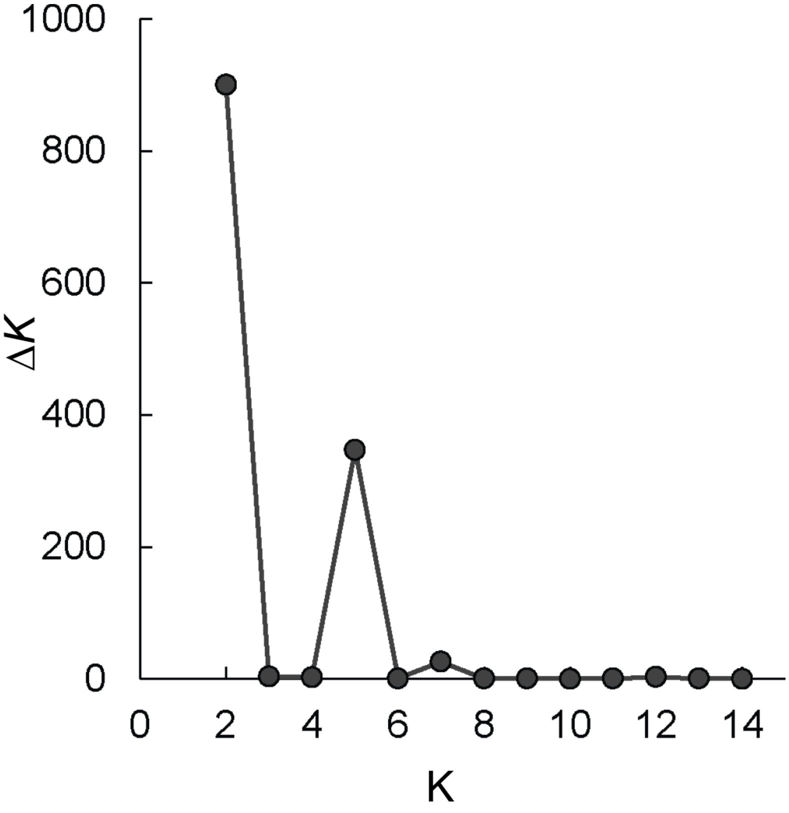


**Supplementary Fig. 10. Estimated Δ*K* values with *K* from 2 to 14.**

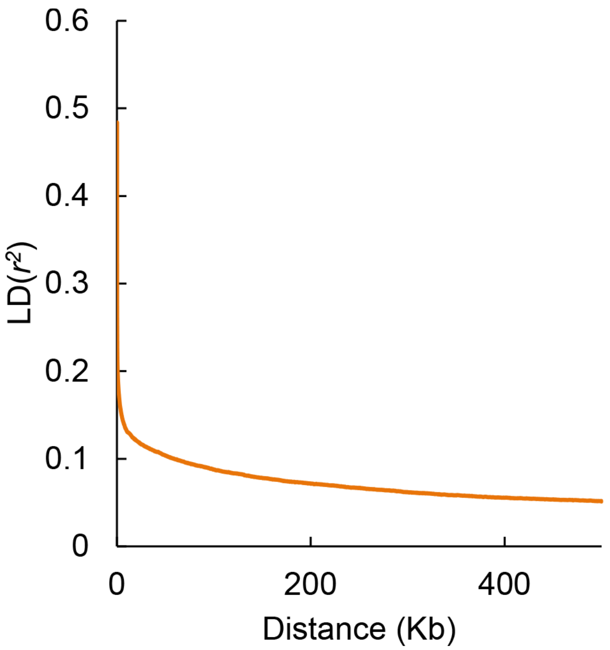


**Supplementary Fig. 11 Linkage disequilibrium (LD) decay pattern of cultivated spinach (*Spinacia oleracea*).**

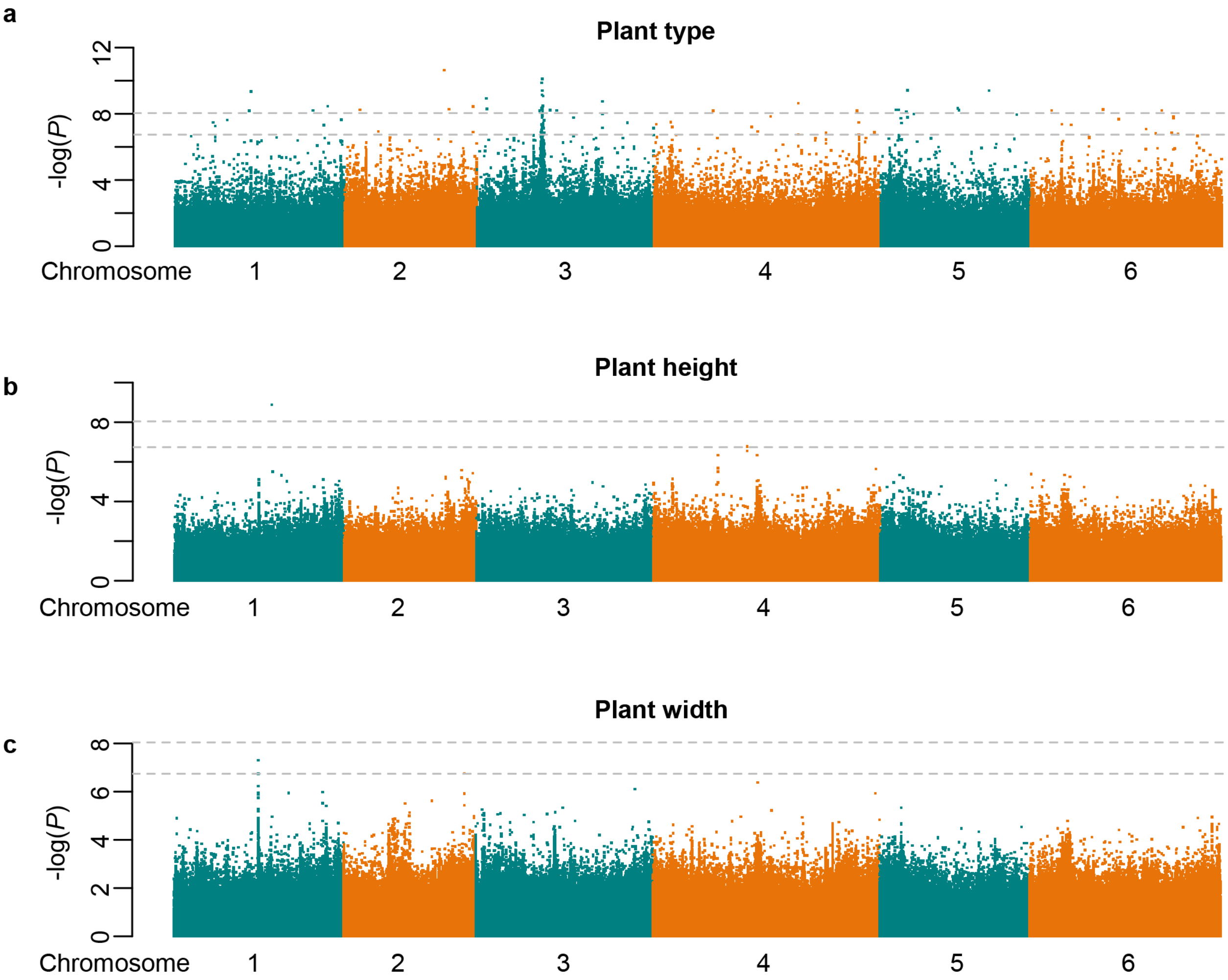


**Supplementary Fig. 12 Manhattan plots of GWAS of plant type (a), plant height (b) and plant width (c).** Gray horizontal dashed lines indicate the Bonferroni-corrected significance thresholds of GWAS ($\alpha = 0.05$ and $\alpha = 1$, respectively).

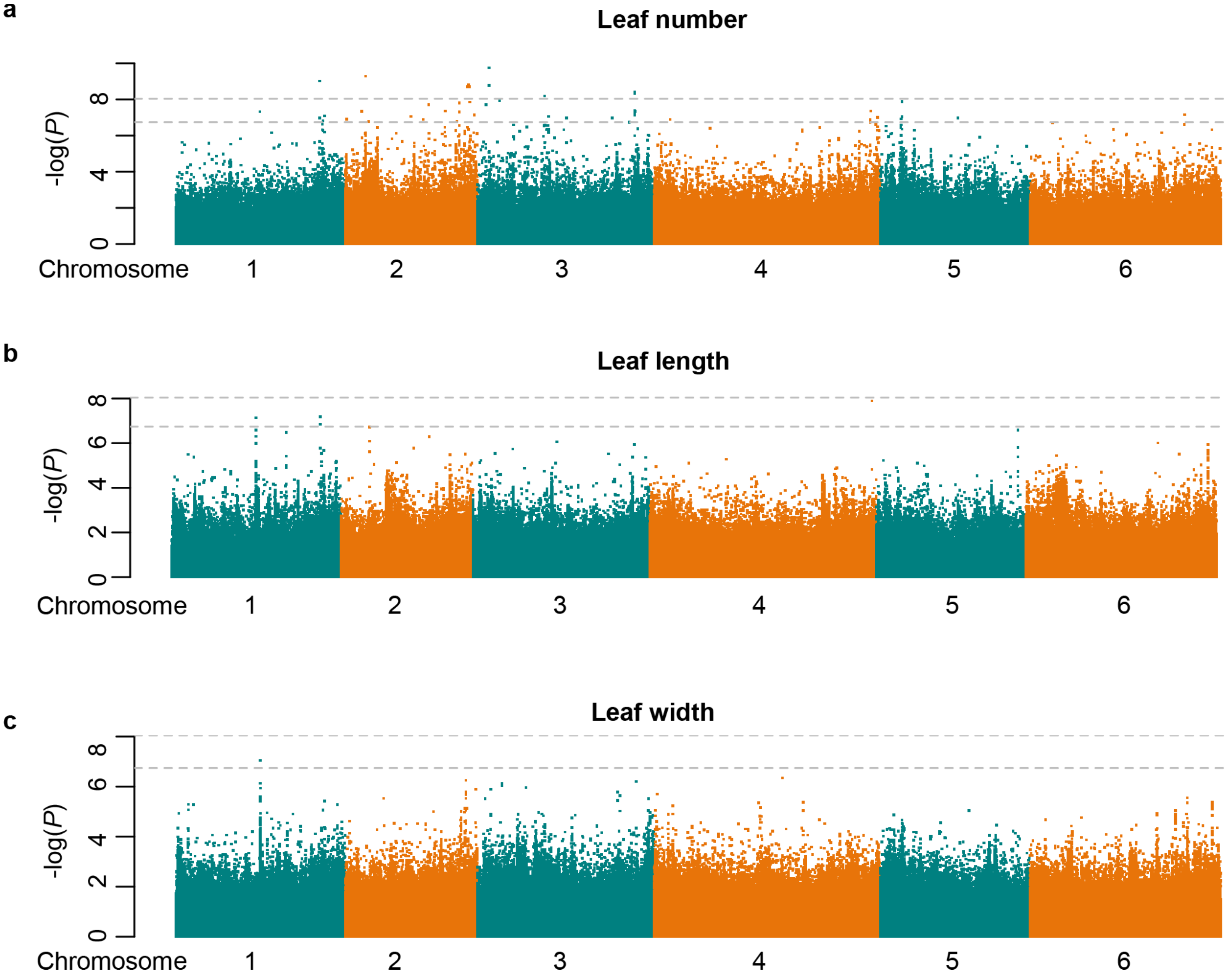


**Supplementary Fig. 13 Manhattan plots of GWAS of leaf number(a), leaf length (b) and leaf width (c).** Gray horizontal dashed lines indicate the Bonferroni-corrected significance thresholds of GWAS ($\alpha = 0.05$ and $\alpha = 1$, respectively).

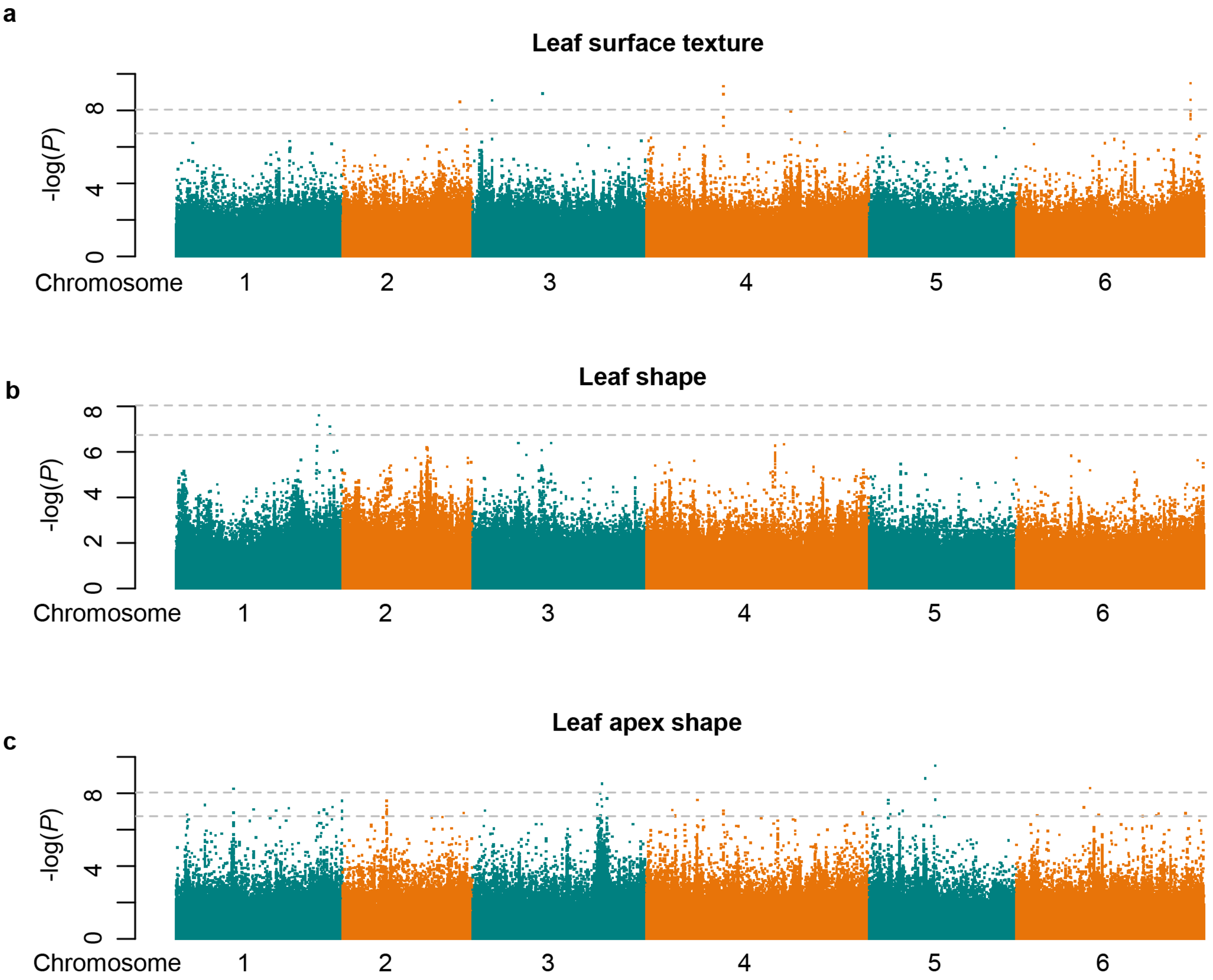


**Supplementary Fig. 14 Manhattan plots of GWAS of leaf surface texture (a), leaf shape (b) and leaf apex shape (c).** Gray horizontal dashed lines indicate the Bonferroni-corrected significance thresholds of GWAS (α = 0.05 and α = 1, respectively).

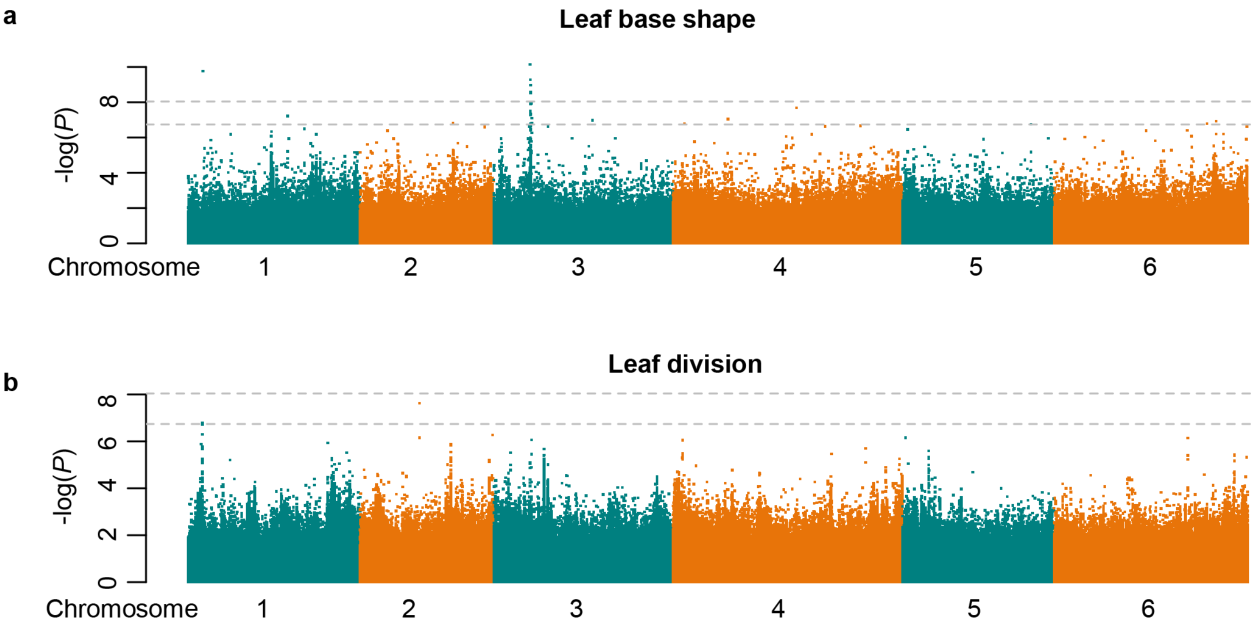


**Supplementary Fig. 15 Manhattan plots of GWAS of leaf base shape (a) and leaf division (b).** Gray horizontal dashed lines indicate the Bonferroni-corrected significance thresholds of GWAS (α = 0.05 and α = 1, respectively).

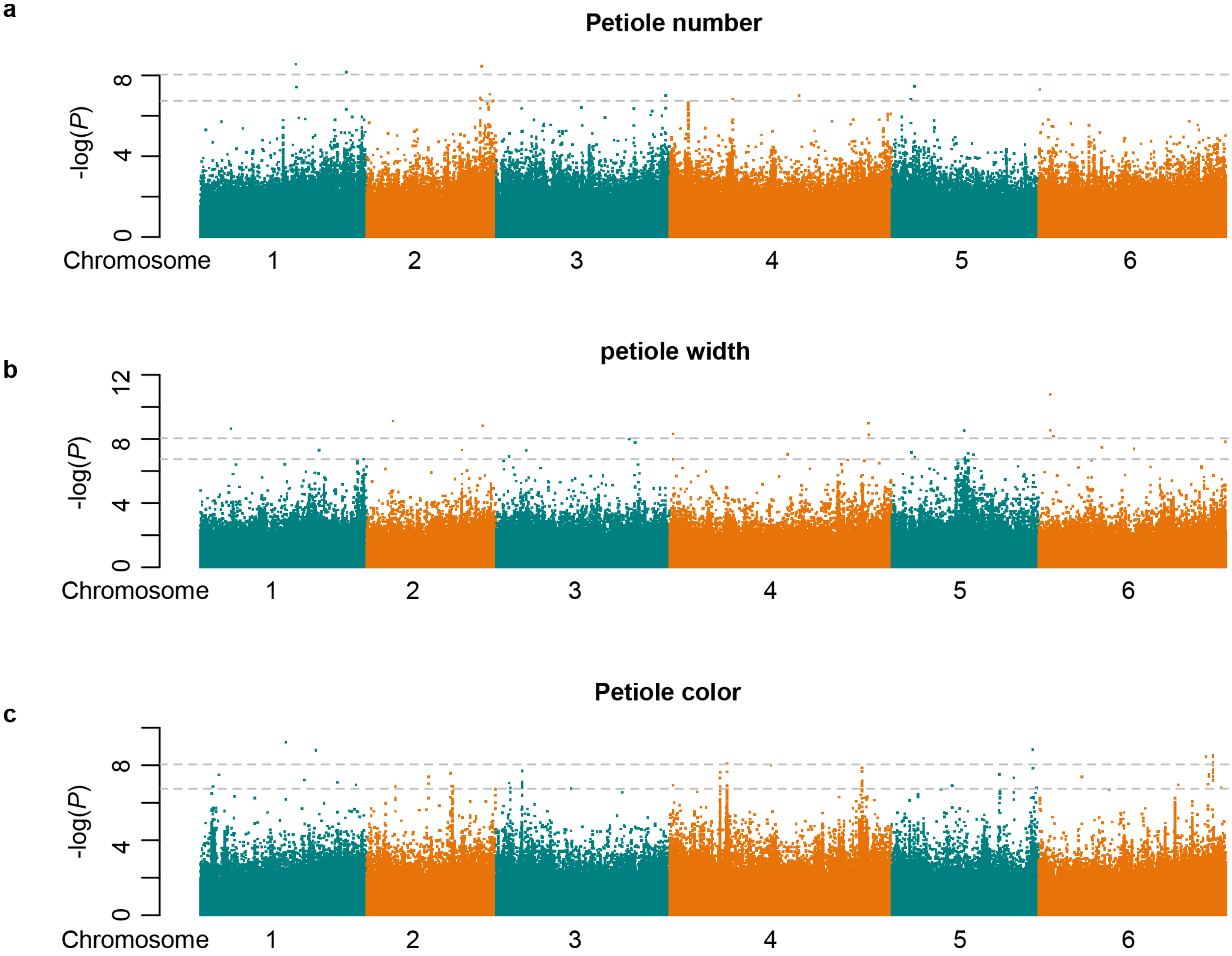


**Supplementary Fig. 16 Manhattan plots of GWAS of petiole length (a), petiole width (b) and petiole color (c).** Gray horizontal dashed lines indicate the Bonferroni-corrected significance thresholds of GWAS (α = 0.05 and α = 1, respectively).

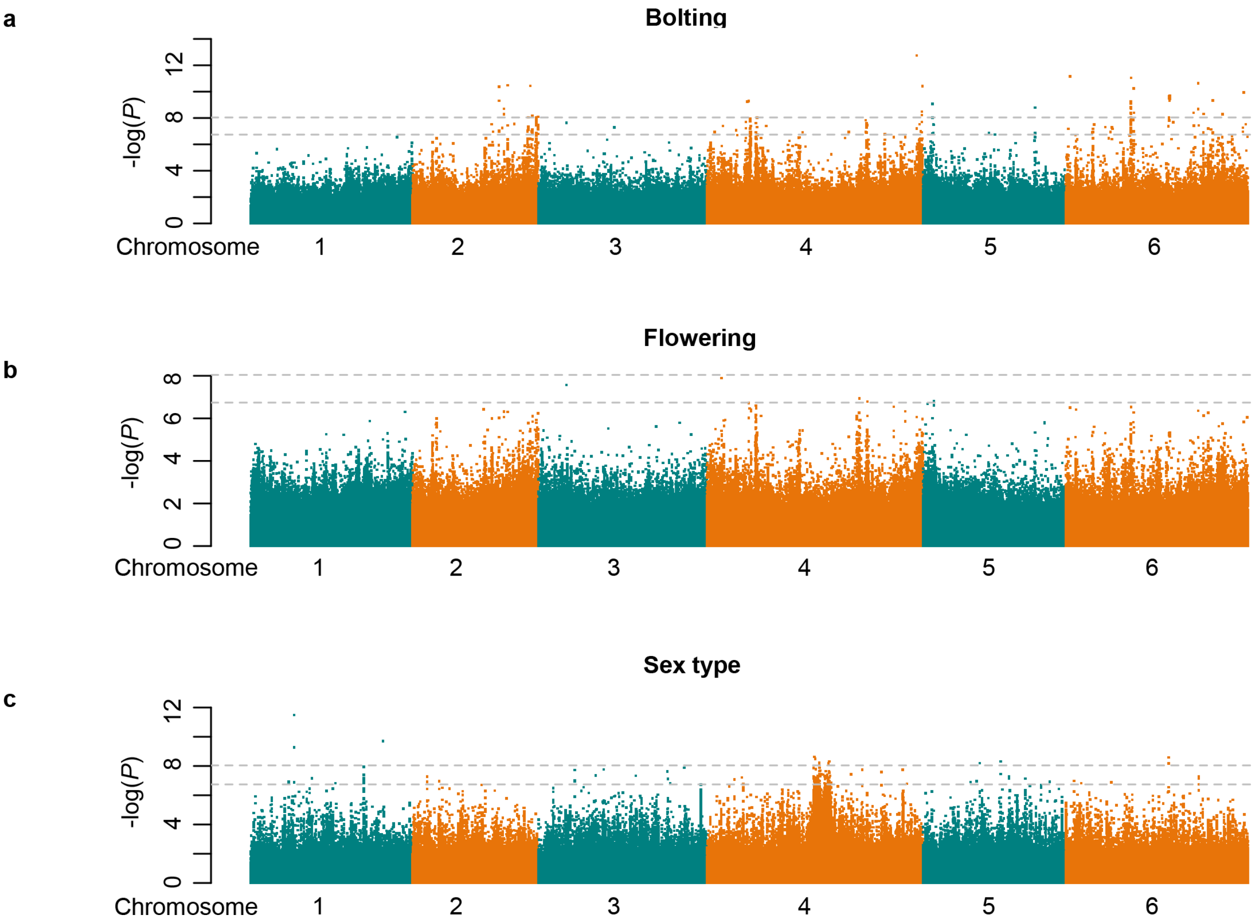


**Supplementary Fig. 17 Manhattan plots of GWAS of bolting (a), flowering (b) and sex type (c).** Gray horizontal dashed lines indicate the Bonferroni-corrected significance thresholds of GWAS (α = 0.05 and α = 1, respectively).

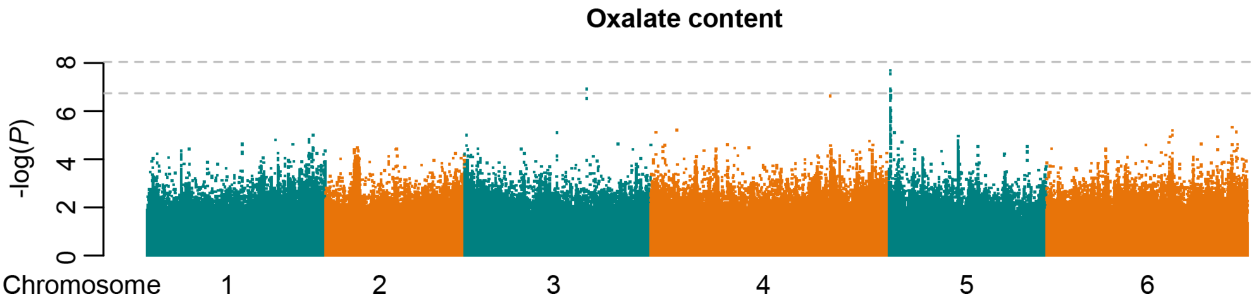


**Supplementary Fig. 18 Manhattan plots of GWAS of oxalate content.** Gray horizontal dashed lines indicate the Bonferroni-corrected significance thresholds of GWAS (α = 0.05 and α = 1, respectively).

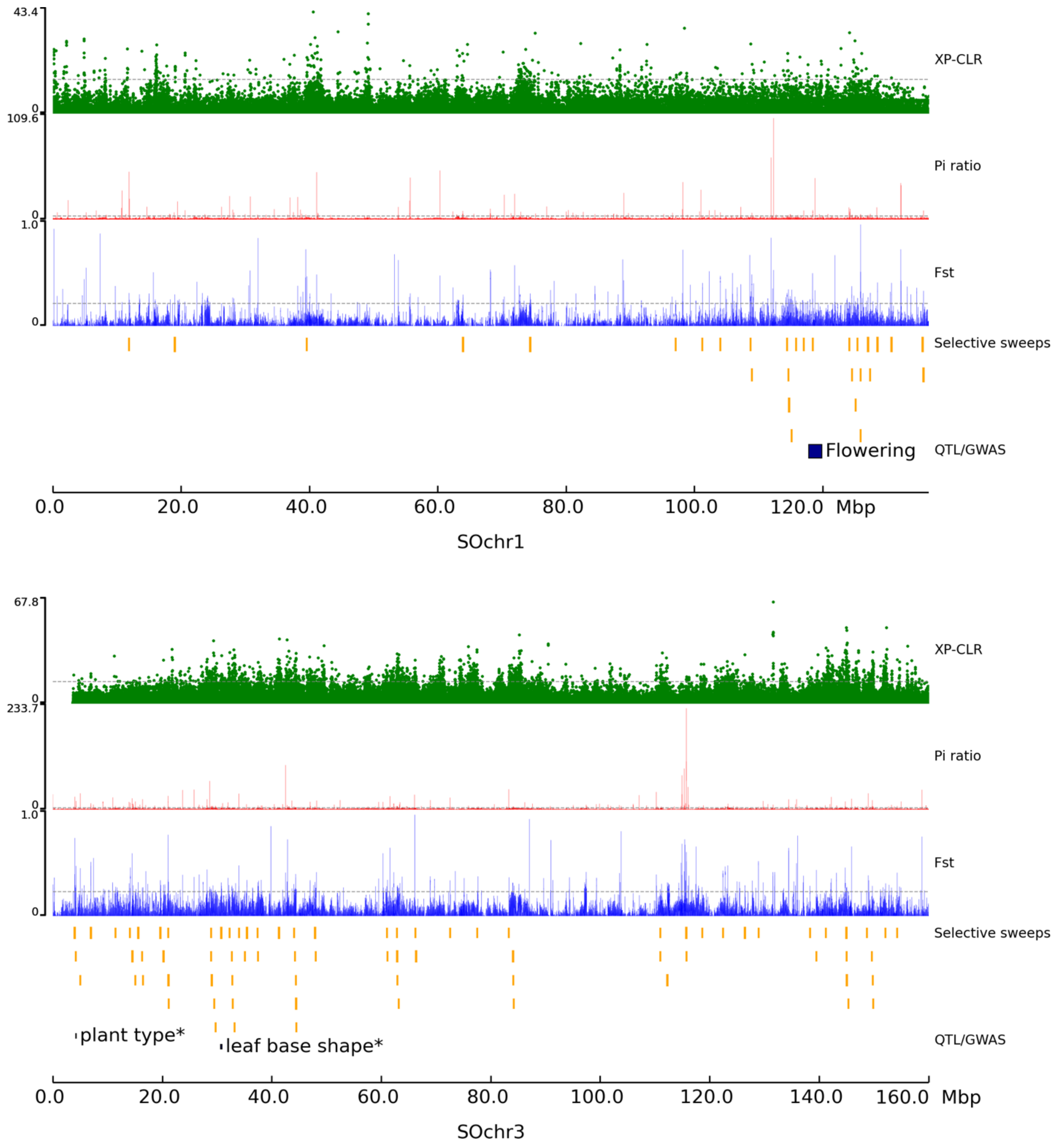


**Supplementary Fig. 19 Genome-wide screening of selective sweeps during spinach domestication.** Gray horizontal dashed lines indicate the top 1% thresholds of the scores/values derived from three approaches, XP-CLR, $F_{ST}$ and nucleotide diversity (π) ratio. Putative selected regions (orange rectangles) and QTL/GWAS signals overlapping with the identified selective sweeps (black rectangles) are shown below these tracks. Traits with * indicate that corresponding GWAS signals were identified in this study.

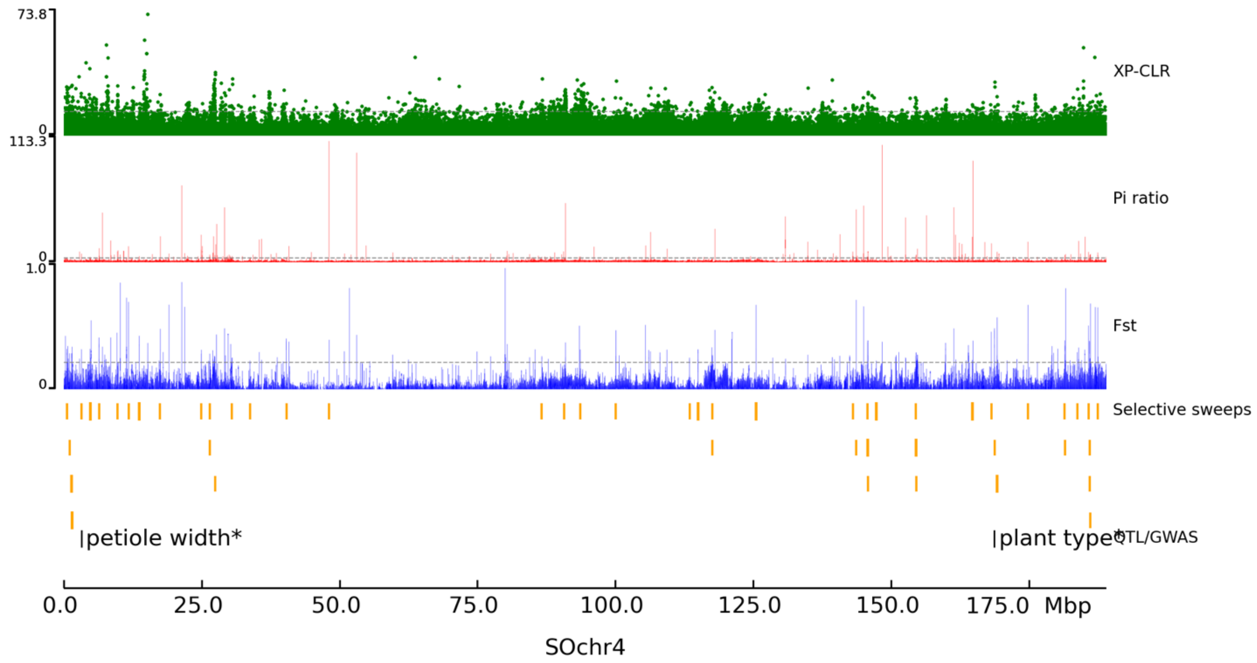


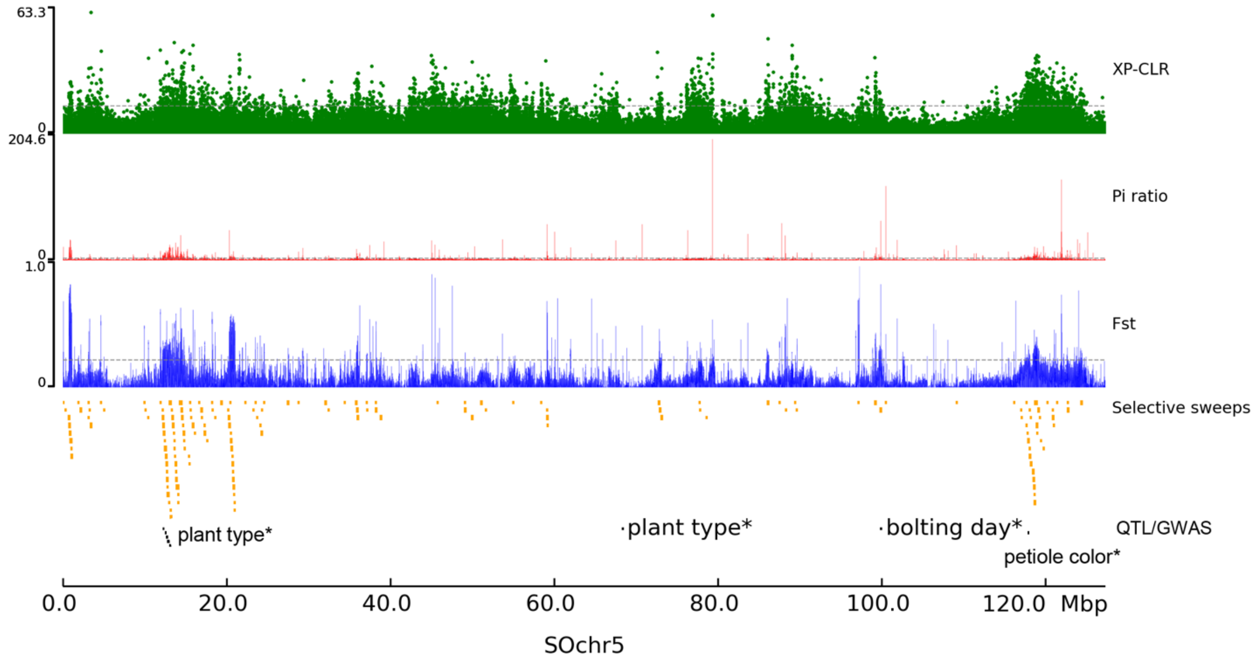


**Supplementary Fig. 19** (continued)

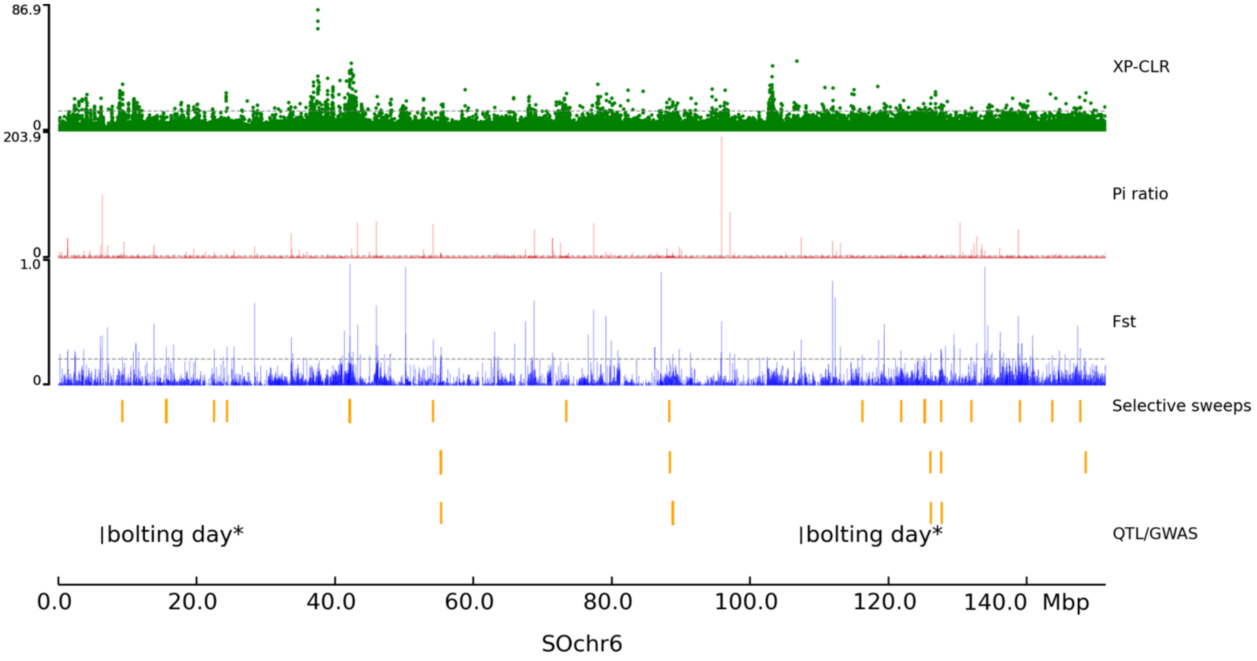


**Supplementary Fig. 19** (continued)
